# Effect of habitual reading direction on saccadic eye movements

**DOI:** 10.1101/2022.01.18.476817

**Authors:** A Lyu, L Abel, AMY Cheong

## Abstract

Cognitive processes can influence the characteristics of saccadic eye movements. Reading habits, including habitual reading direction, also affects cognitive and visuospatial processes, favouring attention to the side where reading begins. Few studies have investigated the effect of habitual reading direction on saccade directionality of low-cognitive-demand stimuli (such as dots). The current study examined horizontal prosaccade, antisaccade and self-paced saccade in subjects with two primary habitual reading directions. We hypothesised that saccades responding to the target in subject’s habitual reading direction would show a longer prosaccade latency and lower antisaccade error rate (errors being a reflexive glance to a sudden-appearing target, rather than a saccade away from it). Sixteen young Chinese participants with primary habitual reading direction from left to right and sixteen young Arabic and Persian participants with primary habitual reading direction from right to left were recruited. Subjects needed to look towards a 5^°^/ 10^°^target in the prosaccade task or look towards the mirror image location of the target in the antisaccade task and look between two 10-degree targets in the self-paced saccade task. Only Arabic and Persian participants showed a shorter and directional prosaccade latency towards 5^°^target against their habitual reading direction. No significant effect of primary reading direction on antisaccade latency towards the correct directions was found. However, we found that Chinese readers generated significantly shorter prosaccade latencies and higher antisaccade directional errors compared with Arabic and Persian readers. The present study provides an insight into the effect of reading habits on saccadic eye movements in response to low-cognitive-demand stimuli and offers a platform for future studies to investigate the relationship between reading habits and neural mechanisms of eye movement behaviours.

## 1. Introduction

### 1.1 Saccadic eye movements

Humans do not look at a scene with steady gaze. Our eyes move around, bringing the interesting parts of the scene to the fovea with the frequency of 2 or 3 fixations per second [1]. In fact, saccadic eye movement is one of the fastest movements produced by the human body, serving in bringing the images of objects of interest into central vision for detailed analysis. The perception of the environment relies on saccades and fixations which are the stops in-between saccades [2]. A distributed network including cortical (mainly frontal and parietal) and subcortical (basal ganglion, superior colliculus, midbrain, brain stem, thalamus and cerebellum) areas are involved in generating saccades [3]. It has been suggested that understanding the saccadic system provides researchers a valuable “microcosm of the brain” as its input can be controlled and manipulated, while its output can be accurately recorded and quantified using different experimental paradigms [4]. A range of eye movement tasks have been used in the literature to examine the characteristics of saccades, including prosaccade, antisaccade and self-paced saccade tasks. Prosaccades, which are also known as reflexive saccades, test the response time (latency) and accuracy of a saccade (saccade gain in terms of the ratio of saccade amplitude / target amplitude) to a sudden-onset peripheral visual stimulus. An antisaccade requires the suppression of a reflexive saccade towards a sudden-onset stimulus and the execution of a voluntary saccade to the opposite direction of the stimulus. The parallel nature of antisaccade programming assumes a competition arises between the exogenously triggered prosaccade and the endogenously initiated antisaccade at the onset of stimulus [5-7]. For example, if the exogenously triggered prosaccade is programmed too fast (or the endogenously initiated antisaccde is too slow to reach the threshold for activation), it “wins” the competition and make a reflexive saccade first (i.e., antisaccade directional error), followed by a corrective antisaccade [8]. Directional error rate (i.e., the proportion of glances towards the stimulus) and the latency of correct response are commonly analysed in antisaccade task. Self-paced saccade task has been considered as an almost entirely volitional eye movement task that requires repetitive and self-initiated refixations between two static visual stimuli [9].

Poor performance in saccadic eye movement has been demonstrated in various neurological and psychiatric disorders [10], such as schizophrenia [11], attention-deficit hyperactivity disorder (ADHD) [12-14], Parkinson’s disease [3, 15, 16] and depression [17]. In particular, a vast array of studies have suggested that many cognitive processes, including those involved in attention [18-20], working memory [21] and learning [22], have an impact on the characteristics of saccadic eye movements.

### 1.2 Effect of cognitive process on saccadic eye movement

Attention is needed to orient the target location prior to the execution of a saccade [20]. Saslow reported a decrease in prosaccade latency from 200 msec to 150 msec if the stimulus appeared 200 msec or longer after the termination of the fixation, compared to the situation where the offset of fixation and onset of stimulation occurred simultaneously [23]. By introducing a medium temporal gap (200 – 250 msec) between the offset of a central fixation target and the onset of a peripheral stimulus, Fischer and Weber found a significant decrease in antisaccade latency but a significant increase in antisaccade error rate [18]. One explanation for these changes in saccades was that this temporal gap contributed to the disengagement of attention before the target appeared. Moreover, studies manipulating the likelihood of the target presenting on either left or right side of a central fixation point found that subjects showed shorter prosaccade latency to the target direction with a higher probability of presentation. This suggested an effect of learning in modifying the prosaccade performance [22, 24].

While changing the direction of letters and words within English sentences (i.e., both letters and words were orientated from right to left), Inhoff et al. reported less efficient saccadic eye movements in English readers compared with their reading normal English texts. However, these performances improved with practice [25]. In addition to these studies, extensive findings have revealed a wide range of cognitive processes influencing saccadic eye movements [18-20]. Even a simple prosaccade involves a complex weighting of both bottom-up information (stimulus properties) and top-down information (cognitive factors), although the precise nature for the degrees of the control remains unclear [8]. In addition, our cognitive systems are shaped or influenced by cultural practices such as reading habits (see below of section 1.3), which suggests the impact of reading habits on characteristics of saccadic eye movements.

### 1.3 Effect of habitual reading direction on cognitive systems

Han and Northoff (2008a) provided neuroimaging evidence that transcultural differences could affect the neural activities underlying both high-level and low-level cognitive functions [26]. They proposed to investigate the influence of reading direction on regulating the functional organization of the human brain as well as related neurocognitive processes [27]. Reading direction has been found to influence many cognitive functions, such as directional differences in facial expression perception [28], aesthetic preference [29] and utilization of visual space [30]. Especially, visuospatial attention can be modulated by the habitual reading direction. During a letter matching task, English readers with habitual reading direction of left-to-right (LTR) spent a longer time in responding to the stimulus in the right visual field, while Hebrew readers with habitual reading direction of right-to-left (RTL) took longer to respond to the stimulus that appeared in the left visual field. They suggested that reflexive attention showed biases on the side where reading began [31]. Consistent with this study, several studies have confirmed the effect of habitual reading direction on the asymmetries of visuospatial attention [32-34]. For example, Rinaldi and colleagues (2014) compared the performance on a star cancellation task between Italian and Israeli subjects who were instructed to mark the small stars amongst many randomly distributed distractors (large stars, English or Hebrew letters and words). They found that monolingual Italian subjects (i.e., reading from LTR) made more omissions in the right visual field, while monolingual Israeli subjects (i.e., reading from RTL) omitted more targets in the left visual field [33]. However, bilingual subjects who managed reading in both directions did not show any spatial asymmetries. Further, Afsari and colleagues (2016) examined the effect of habitual reading direction in bilingual readers of a native LTR language and a secondary RTL language. They found that native LTR readers who studied a secondary RTL language in late life showed a leftward bias with more fixations on the left part of a natural image, and this horizontal bias of the exploration of images did not alter when they first read either LTR or RTL text primes [34].

In addition to the biased visuospatial attention, we questioned whether reading direction also contributed to the left-right asymmetry of saccadic eye movements. While reading continuous text, LTR texts such as English [35] and German [36] elicit saccades towards the location slightly to the left of a word centre, while RTL scripts such as Hebrew [37] and Uighur [38] have saccades landing to the position slightly to the right of a word centre. In addition to saccades generated towards to the left and right directions, Yan et al. took the advantage that Chinese text can be orientated horizontally and vertically without disturbing the shape of characters and reported a similar saccadic landing position for 28 young readers reading horizontal and vertical Chinese texts [39]. These studies demonstrated that reading direction affected saccadic eye movements during high-level reading processes. Nevertheless, fewer studies have investigated the impact of the habitual reading direction on the directionality of saccadic eye movements during low-cognitively demanded tasks such as responding to a dot. Most of the studies investigating the left-right asymmetry of these saccadic eye movements focused on ocular dominance [40-42] or hand dominance [43, 44]. Understanding the effect of habitual reading direction on saccadic eye movements of low-cognitive-demand stimuli would help researchers to better investigate the differences of eye movement control across populations.

In the present study, young healthy participants with two primary habitual reading directions were recruited to complete 3 types of saccadic eye movement tasks, namely horizontal prosaccade, antisaccade and self-paced saccade. These encompassed both reflexively and volitionally initiated saccades. We hypothesized that readers who habitually read from LTR should show a leftward asymmetry in the saccadic parameters (i.e., shorter prosaccade latency and higher antisaccade error rate) when they made a saccade responding to the target appeared in the direction where reading began (i.e., the target appeared at the left of a fixation point) compared with the target appeared along their habitual reading direction (i.e., the target appeared at the right of the fixation point). In contrast, readers who habitually read from RTL would show a rightward asymmetry.

## 2. Material and methods

### 2.1 Subjects

32 young university students who were bilingual speakers and readers were recruited from the University of Melbourne (20) and The Hong Kong Polytechnic University (16). The calculated sample size provided 85% power to detect a significant difference (an estimated effect size of 0.71) between the 2 reading directions of 5^°^stimulus at the one-tailed 0.05 alpha level. 16 subjects were Chinese readers whose primary reading direction was LTR and 16 Arabic and Persian readers (12 Arabic and 4 Persian) whose primary reading direction was RTL. All participants were aged between 18 and 35 with normal or corrected to normal vision and started to learn English (with reading direction of LTR) since their early childhoods. To control the potential confounding influence of education level, this factor was controlled and matched between these 2 groups. Participants in the RTL group were significantly older than the LTR group. Nevertheless, horizontal saccade latency is relatively stable from age of 14 to 50 years old [45], so this factor should not be a concern. Exclusion criteria were any history of ophthalmic, neurological or psychotic illness, or any medications intake that might affect eye movements. Subjects were separated into 2 groups according to their primary habitual reading direction. The characteristics of the participants are shown in Table 1. Informed consent was obtained in accordance with a protocol approved by the University of Melbourne human research ethics committee (HREC #1647981.1) and Department of Research Committee of the School of Optometry of The Hong Kong Polytechnic University (HSEARS20191217001). The study followed the tenets of the Declaration of Helsinki.

**Table 1.**
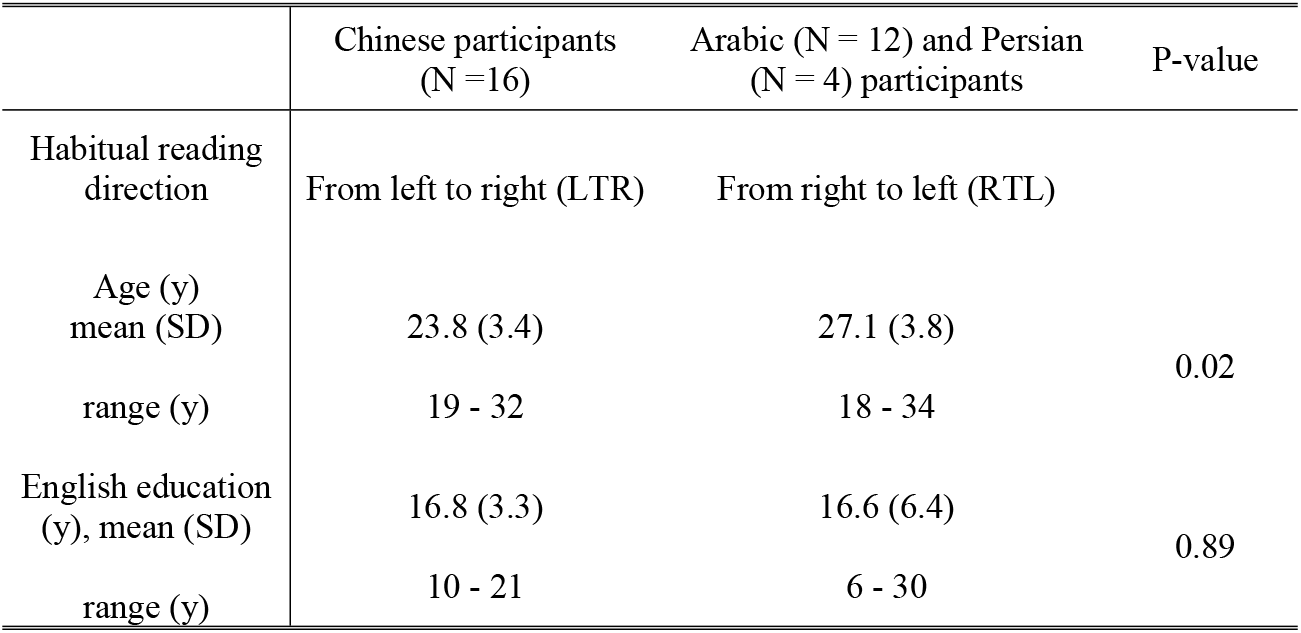
Descriptive characteristics of participants.

|  | Chinese participants<br>(N = 16) | Arabic (N = 12) and Persian<br>(N = 4) participants | P-value |
| --- | --- | --- | --- |
| Habitual reading direction | From left to right (LTR) | From right to left (RTL) |  |
| Age (y)<br>mean (SD) | 23.8 (3.4) | 27.1 (3.8) | 0.02 |
| range (y) | 19 - 32 | 18 - 34 |  |
| English education<br>(y), mean (SD) | 16.8 (3.3) | 16.6 (6.4) | 0.89 |
| range (y) | 10 - 21 | 6 - 30 |  |

### 2.2 Apparatus and stimuli

As the data were collected at 2 sites, minor differences of experimental setting were present, including presenting monitor and testing distance, whereas the stimulus size and distance were adjusted so that same visual angle was elicited. Both the testing sequence and program (including resolution and refresh rate) were identical between the 2 sites. The stimulus was a 1-degree black dot against a white background on a 27-inch (U2711B, Dell Technologies, Round Rock, Texas, United States) or a 24-inch LCD monitor (BENQ xl2540) in Melbourne and Hong Kong site respectively. Participants sat comfortably at 75 cm (Melbourne) or 65 cm (Hong Kong) in front of the monitor with chin resting on a chinrest to stabilize their head position. Movement of both eyes was recorded using an infrared video eye tracking system (Eyelink 1000 or Eyelink Portable Duo, SR Research, Scarborough, ONT, Canada) with a sampling rate of 500 Hz. The resolution and refresh rate of the monitors were 1920 × 1080 pixels and 60 Hz. Subjects were asked to perform the following eye movement tasks.

### 2.3 Procedures

Participants’ eye movements were assessed while conducting 3 visual tasks: 1) prosaccade, 2) antisaccade and 3) self-paced saccade tasks.

Targets were presented pseudorandomly in locations 5 or 10 degrees to the left or right of the centre of the monitor. Participants were instructed to fixate at a centre cross and then to look towards the target in the prosaccade task (see Fig. 1a) or look towards the mirror image of the target in the antisaccade task (see Fig. 1b) as soon as the target was presented and fixation disappeared. Fifty-two trials were conducted in the prosaccade task to assess the prosaccade latency (i.e., reaction time responding to the onset of stimulus) and gain (i.e., ratio of saccadic amplitude to target amplitude). Express saccades whose latency falls between 80 to 120 msec [46-48] were excluded from the analysis. Less than 15 % of the trials were excluded for all participants. Fifty-two trials were conducted in the antisaccade task to assess the antisaccade latency of correct responses (i.e., saccades made to the correct direction) and error rate (i.e., proportion of prosaccade errors). In the self-paced saccade task, two targets were shown for 45 seconds at 10 degrees left and right of the centre of the monitor. Participants needed to look back and forth between these two dots as rapidly and as accurately as possible for the entire duration of the task. Gain (i.e., ratio between the primary saccade and target amplitude) and inter-saccadic intervals (i.e., interval between onset of the saccades) were collected and submitted to data analysis.

**Fig 1.**
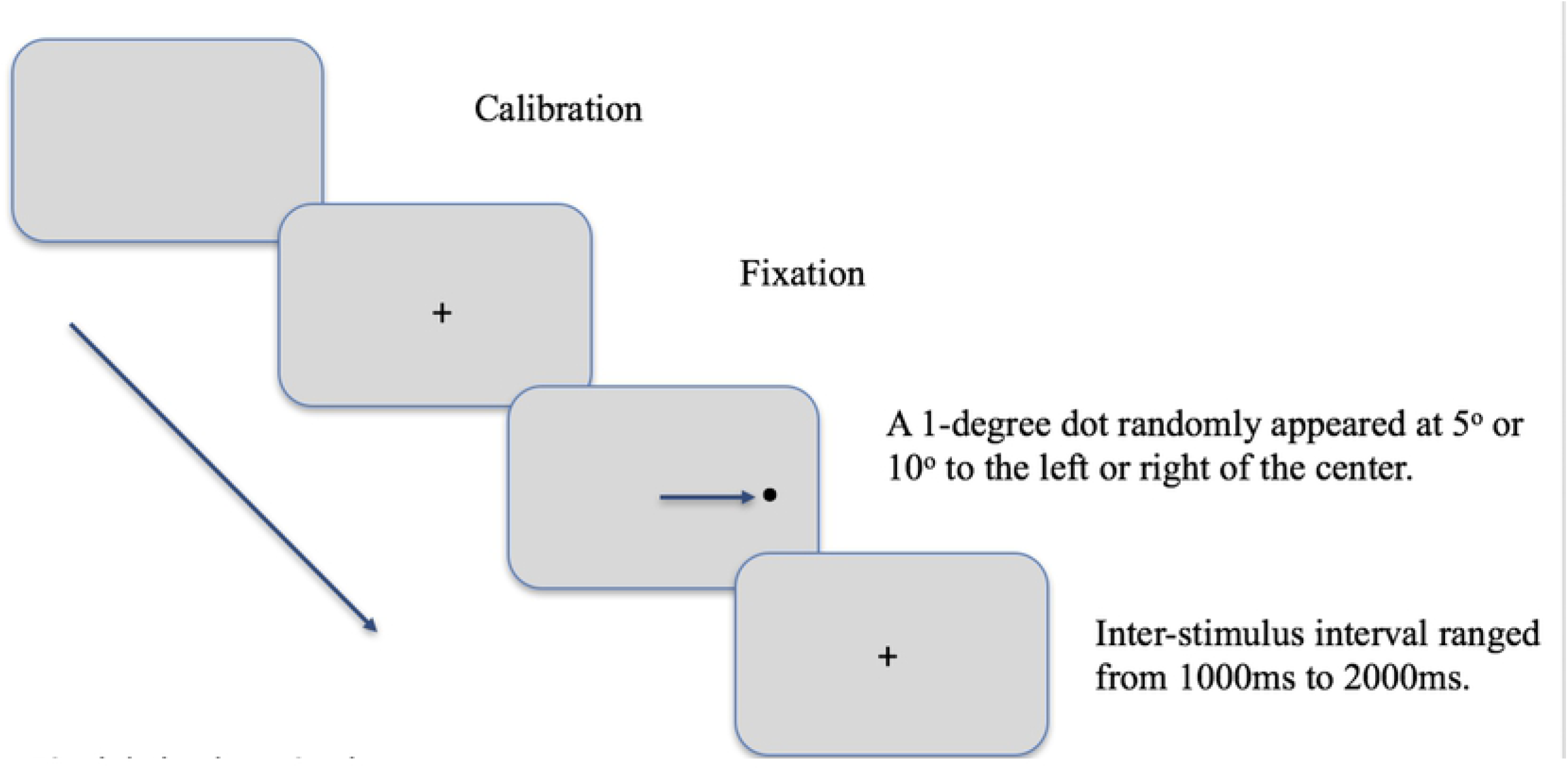
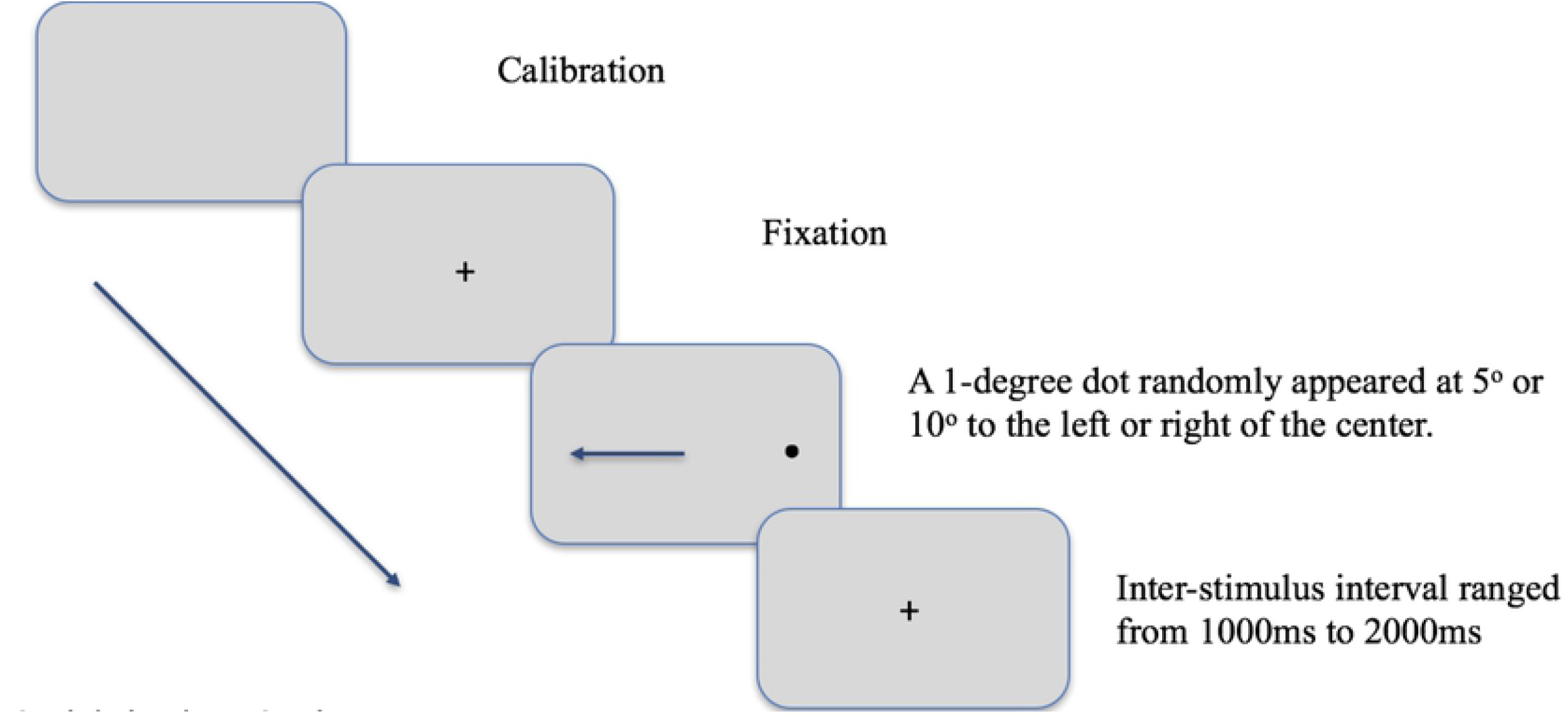
Sample trial of prosaccade and antisaccade task. Fig. 1a. A sample trial of prosaccade task when participants need to make a saccade towards the target as quickly as possible. Fig. 1b. A sample trial of antisaccade task when participants need to look at the mirror image of the target location as quickly as possible.

### 2.4 Data analysis

All statistical analysis was performed using GraphPad Prism version 9.2.0.332 for Windows (GraphPad Software, San Diego, California USA, www.graphpad.com). Eye movement parameters were not significantly different from normal distribution (Kolmogorov-Smirnov goodness of fit test, p > 0.05). Dependent variables (prosaccade latency, prosaccade gain, correct antisaccade latency, antisaccade error rate, inter-saccadic interval, and gain in self-paced saccades) were analysed using analysis of variance (ANOVA) with group (LTR (Chinese participants) vs. RTL (Arabic and Persian participants)) as between-subject factors and the direction of stimulus (with-vs. against habitual reading direction) and / or the magnitude of stimulus from the fixation (5° vs. 10°) as within-subject factors, to assess any significant effect or interaction. A p-value of less than 0.05 was considered statistically significant.

## 3. Results

### 3.1 Effects of habitual reading direction on prosaccade eye movements

The average prosaccade latency of the RTL group in response to a sudden-onset stimulus was significantly longer than that of the LTR group (mean 188.06 vs. 174.41 msec, F(1,60)=5.61, p=0.02). Bonferroni’s post-hoc analysis demonstrated that RTL participants required longer prosaccade latency in responding to targets presented at 5^°^along their habitual reading direction compared with LTR participants (198.98 vs. 167.75 msec, p=0.01). Neither stimulus direction (i.e., presented with-or against-reading direction) or magnitude (i.e., presented at 5° or 10° away from fixation) had a significant effect on prosaccade latency (F(1,60)<1.59, p>0.21), whereas a significant interaction between group and stimulus direction was observed (F(1,60) = 8.07, p=0.006). The RTL participants had longer prosaccade latency when the target was presented 5^°^along their habitual reading direction (i.e., target at the left side of the fixation) compared with that appeared against the reading direction (198.98 vs. 178.77 msec, p=0.03). However, LTR participants showed similar reaction time for the target appearing towards the two directions (167.75 vs. 176.56 msec, p>0.99; Table 2).

**Table 2.** Eye movement characteristics in the prosaccade and antisaccade task (mean and standard deviation)

| Group |  |  |  | Chinese participants<br>(N = 16) (LTR) | Arabic (N = 12) and Persian<br>(N = 4) participants (RTL) | P-value |
| --- | --- | --- | --- | --- | --- | --- |
| Saccade type |  | Stimulus magnitude | Stimulus direction |  |  |  |
| Prosaccade | Prosaccade<br>latency<br>(msec) | 5° | With-direction | **167.75 (22.06)** | **198.98 (31.76)** | **0.01\*** |
|  |  |  | Against-direction | 176.56 (24.64) | **178.77 (23.86)** | >0.99 |
|  |  | P-value |  | >0.99 | **0.03\*** |  |
|  |  | 10° | With-direction | 176.00 (17.91) | 190.33 (29.32) | >0.99 |
|  |  |  | Against-direction | 177.36 (36.82) | 184.15 (19.11) | >0.99 |
|  |  | P-value |  | >0.99 | >0.99 |  |
|  | Prosaccade<br>gain | 5° | With-direction | 1.02 (0.12) | 0.96 (0.10) | >0.99 |
|  |  |  | Against-direction | 0.98 (0.08) | 1.03 (0.23) | >0.99 |
|  |  | P-value |  | >0.99 | 0.16 |  |
|  |  | 10° | With-direction | 0.99 (0.08) | 0.94 (0.08) | >0.99 |
|  |  |  | Against-direction | 0.97 (0.05) | 0.95 (0.11) | >0.99 |
|  |  | P-value |  | >0.99 | >0.99 |  |
| Antisaccade | Antisaccade<br>latency<br>(msec) | 5° | With-direction | 285.73 (45.18) | 277.74 (46.70) | >0.99 |
|  |  |  | Against-direction | 287.11 (38.79) | 276.90 (45.08) | >0.99 |
|  |  | P-value |  | >0.99 | >0.99 |  |
|  |  | 10° | With-direction | 285.96 (30.69) | 273.05 (51.56) | >0.99 |
|  |  |  | Against-direction | 280.21 (33.69) | 275.84 (45.50) | >0.99 |
|  |  | P-value |  | >0.99 | >0.99 |  |

\*: $p < 0.05$

Prosaccade gain examines the accuracy of the landing position of prosaccade. Neither of the three independent variables (i.e., group, stimulus direction and stimulus magnitude) significantly affected the gain (F(1,60)<2.15, p>0.15). Nevertheless, similar to the latency result, a significant interaction between group and stimulus direction was found (F(1,60)=6.93, p=0.01). Although no significance was found in the post-hoc analysis (Table 2).

### 3.2 Effects of habitual reading direction on antisaccade eye movements

Opposite to the findings in prosaccade eye movements, there was no significant main effect or interaction effect of group, stimulus direction and stimulus magnitude on the antisaccade latency for the correct trials (F(1,60)<0.82, p>0.37; Table 2).

When comparing antisaccade errors between different groups and among different stimulus positions, stimulus magnitude (i.e., 5^°^vs. 10°) was found to significantly affect the rate of directional errors (F(1,60)=9.25, p=0.004), that both groups made more antisaccade errors towards 5^°^targets compared with 10^°^targets (rate of 0.24 vs. 0.17 and 0.24 vs. 0.10 for the LTR and RTL group respectively; Fig. 2). A significant interaction between group and stimulus direction (i.e., with-vs. against-reading direction) was observed (F(1,60)=6.92, p=0.01), that the LTR group marginally had more antisaccade errors for targets appearing 10^°^away from the centre at their habitual reading side, compared with the RTL group (rate of 0.22 vs. 0.07, p=0.05).

**Fig 2.**
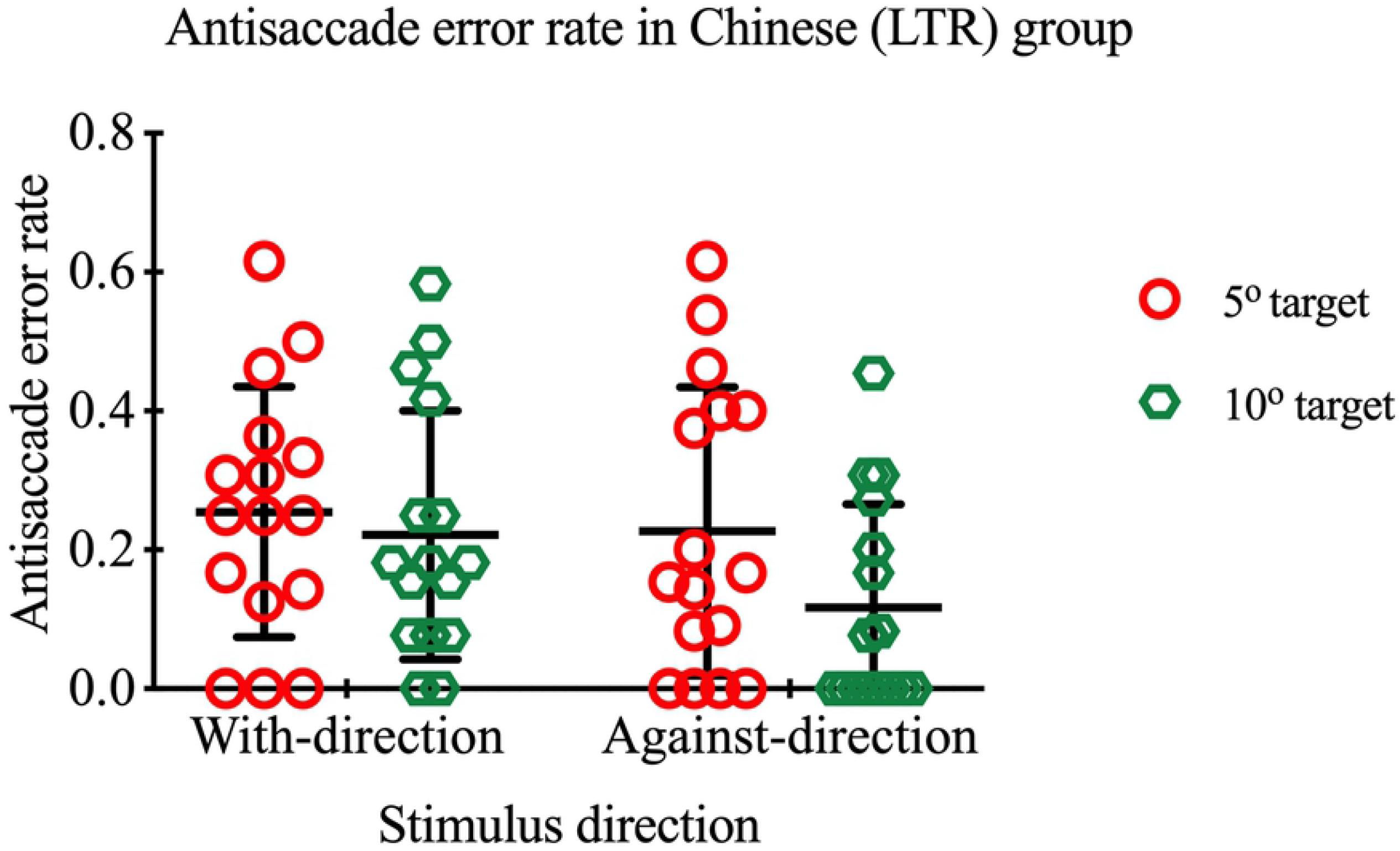
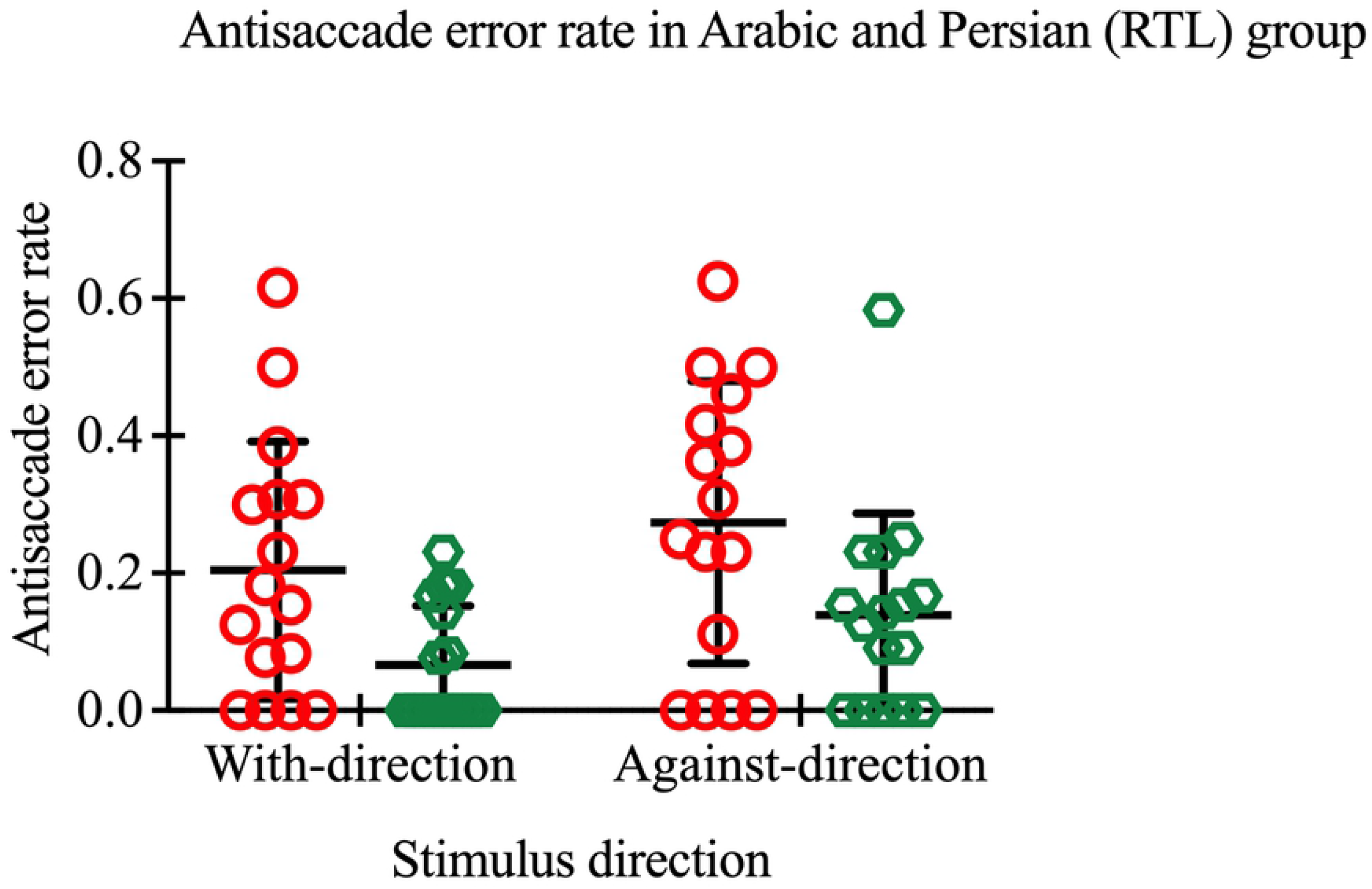
Antisaccade error rate in 2 groups of participants. The antisaccade error rate of the Chinese (left panel) as well as the Arabic and Persian group (right panel) in responding to the 5^°^and 10^°^with-direction target was 0.25 ± 0.18 and 0.22 ± 0.18, and 0.20 ± 0.19 and 0.07 ± 0.09, respectively, and those for against direction target was 0.23 ± 0.21 and 0.12 ± 0.15, and 0.27 ± 0.21 and 0.14 ± 0.15 respectively. Bars are mean value and standard deviation.

### 3.3 Relationship between prosaccade and antisaccade performance

Further analysis was performed to examine the impact of type of saccade (prosaccade vs. antisaccade) on saccade latency. Interestingly, Chinese participants (LTR group) tended to have shorter prosaccade (174.42 vs. 188.06 msec) but longer antisaccade latency (284.75 vs. 275.88 msec) than the Arabic and Persian participants (RTL group), although this did not reach significance (F(1, 60)=3.74, p=0.06) (see Fig. 3).

**Fig 3.**
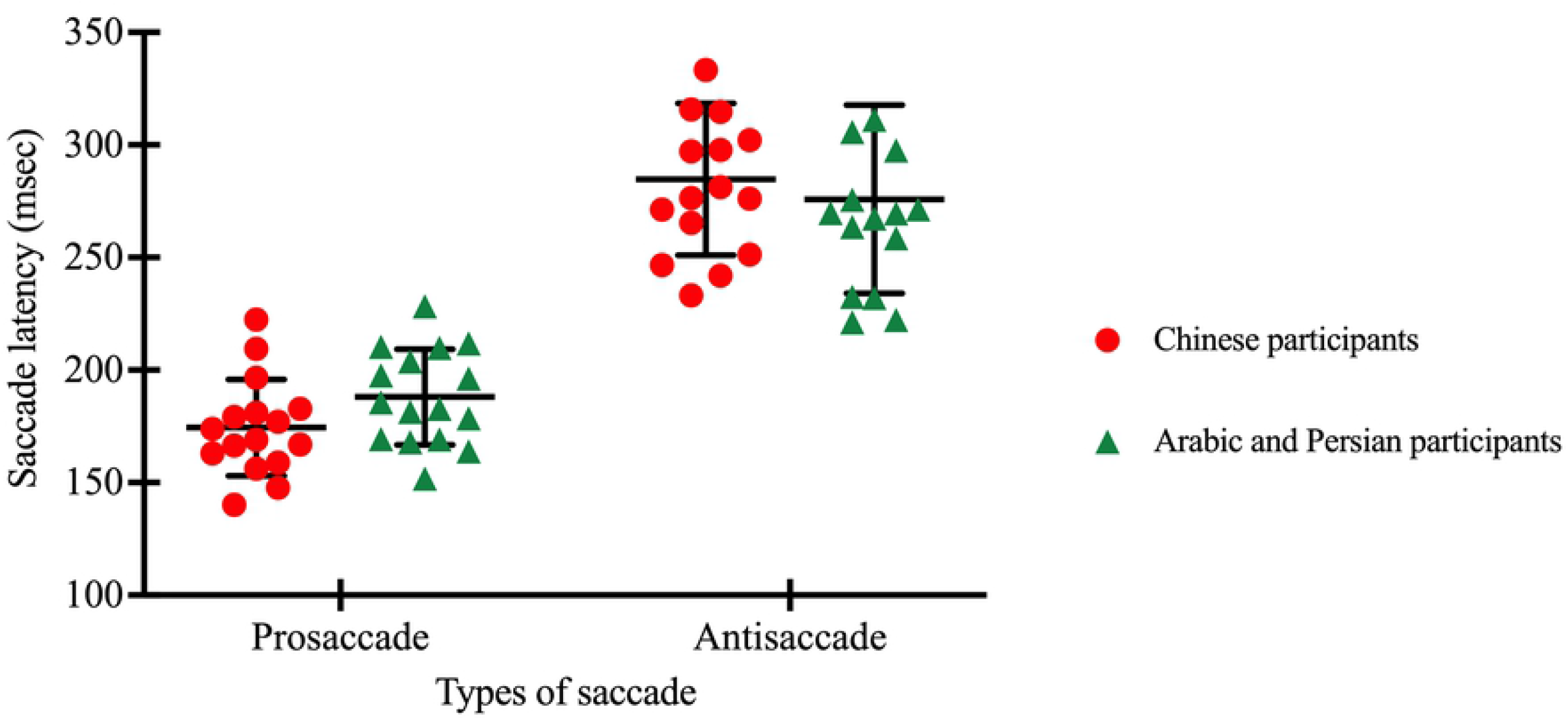
Prosaccade latency and correct antisaccade latency of Chinese (LTR) as well as Arabic and Persian group (RTL) Mean prosaccade latency for the Chinese as well as the Arabic and Persian groups was 174.42 and 188.06 msec respectively, while the mean of correct antisaccade latency was 284.75 and 275.88 msec respectively.

### 3.4 Effects of habitual reading direction on self-paced saccadic eye movements

Inter-saccadic interval and gain was compared between groups and directions of self-paced saccades (i.e., saccades towards habitual reading direction vs. towards non-habitual reading direction) as well as the interaction effect. No significant group or direction effect was found on interval (F(1,30)<0.14, p>0.71; mean of 509.77 vs. 514.57 msec in the Chinese group and 512.55 vs. 514.29 msec in the Arabic and Persian group for saccades made towards habitual and non-habitual reading direction respectively).

However, a significant interaction between group and saccadic direction was observed on self-paced saccade gain (F(1,30)=14.37, p<0.001; see Fig. 4). The Chinese group showed more accurate gain when they made a saccade to the dot located at the side of their habitual reading direction (i.e., the dot at the right of the monitor) compared with the dot located in their non-habitual reading direction (mean 1.01 vs. 0.95, p=0.02). Whereas participants in the Arabic and Persian group generated more accurate saccades towards the dot showing along their non-habitual reading direction (i.e., dot at the right side of the centre of the monitor) (1.01 vs. 0.96, p=0.03).

**Fig 4.**
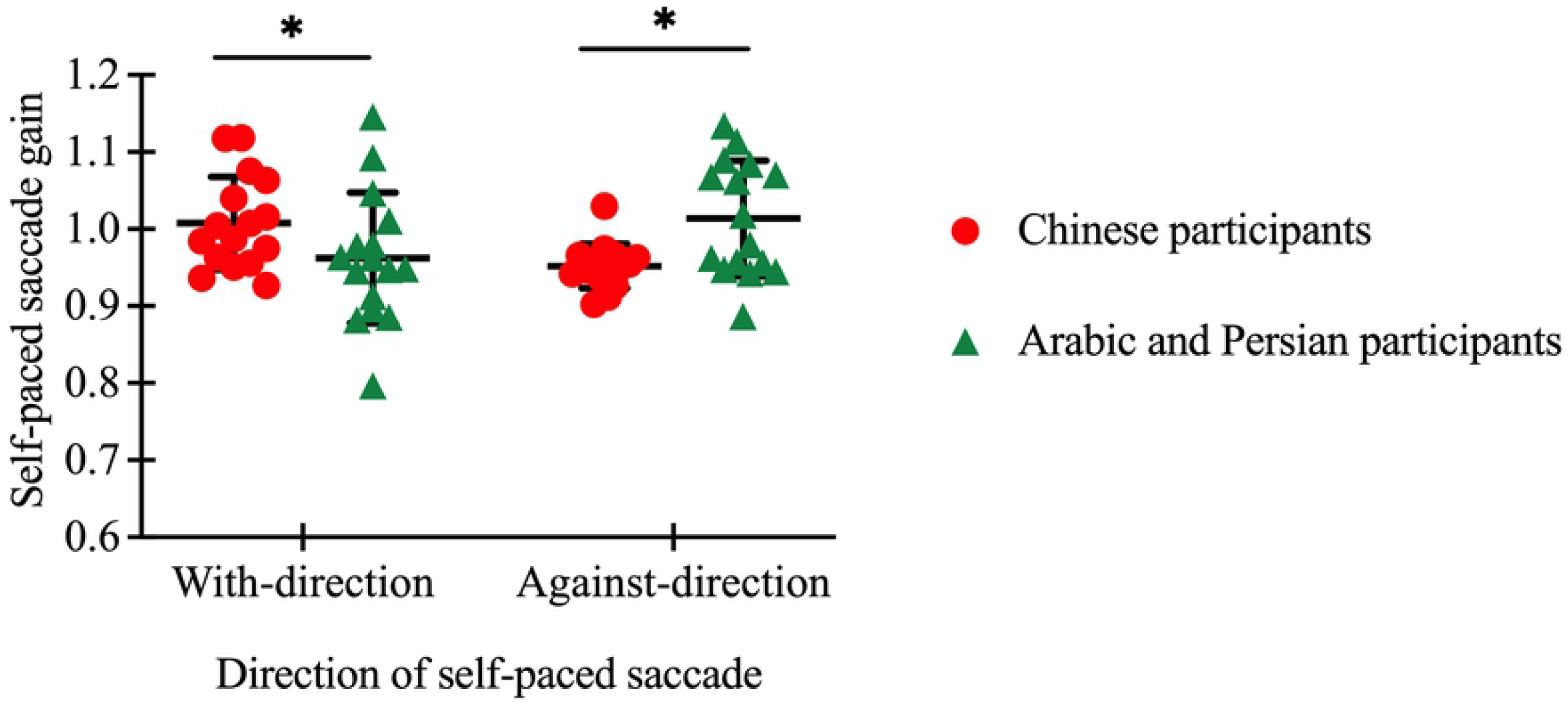
Self-paced saccade gain in Chinese (LTR) as well as the Arabic and Persian group (RTL) The mean of self-paced saccade gain for the Chinese as well as the Arabic and Persian groups made towards the dot showing at their habitual reading direction was 1.01 and 0.96 respectively, while that made towards the dot located at the side of their non-habitual reading direction was 0.95 and 1.01 respectively. *: p<0.05

## 4. Discussion

The objective of this study was to evaluate the impact of the primary habitual reading direction on the directionality of saccadic eye movements to low-cognitive-demand stimuli in young and healthy Chinese as well as Arabic and Persian participants using prosaccade, antisaccade and self-paced tasks. One of the major findings was the significantly shorter saccade latency of the Chinese participants whose primary habitual reading direction was from left to right (LTR) in the prosaccade task compared with that of the Arabic and Persian participants whose primary habitual reading direction was from right to left (RTL). However, the effect of reading direction on the antisaccade latency disappeared, where participants in both groups had similar latencies of accurate antisaccade. The second major finding was that the Chinese subjects generated marginally but significantly more directional errors compared with the Arabic and Persian subjects when the target appeared at 10^°^along their habitual reading direction in the antisaccade task.

### 4.1 Impact of habitual reading direction on prosaccade latency

In this study, we hypothesized that participants would produce shorter prosaccade latency to a stimulus which appeared in their non-habitual reading direction (i.e., left for the Chinese participants and right for the Arabic and Persian participants). This was, however, only found in the Arabic and Persian participants in responding to the 5^°^target. Previous studies reported that the direction of the stimulus presentation did not significantly affect the prosaccade latency of young participants, although previous studies did not consider the participants’ reading direction [41, 49]. In addition, our study found that Chinese readers had 16% shorter prosaccade latency than Arabic and Persian readers when target appeared 5^°^along their habitual reading direction. Amatya et al. reported that Chinese participants generated more low latency ‘express saccades’ compared to non-Chinese participants (Caucasian participants) in an overlap prosaccade task [50]. Their study argued that this difference in saccade latency should be attributed to human genetic diversity rather than cultural differences, as those Chinese participants who grew up in the UK also showed the same pattern of saccade latency as the participants lived in mainland China [51]. A similar study by Knox and colleagues evaluated the antisaccade performance of Chinese participants and found a significantly higher antisaccade directional error rate in those Chinese participants who exhibited a higher proportion of express saccades [52]. They suggested that there was a difference in neurophysiological substrate concerned with eye movement that was not associated with culture. Nevertheless, in addition to express saccade latency, few studies have demonstrated the difference in normal reflexive saccades with both top-down and bottom-up control between populations. An electrophysiological study in primates showed that a neural signal took around 40 msec to be transmitted from the retina to the superior colliculus (SC), and it took approximately 20 msec to stimulate the SC to trigger a saccadic eye movement to a specific location [53]. However, the typical latency of a prosaccade is around 200 msec in humans [8]. Carpenter argued that such a long latency of the saccadic eye movement was due to the decision time on making a decision to look at the target or not [54]. One possible explanation for the different reaction time across groups was that these two groups’ participants used different decision-making strategies, resulting in different decision-making times. However, further study is required to investigate the decision-making time for simple cognitive tasks between different populations.

It has been suggested that attention needs to orient to the target location prior to the execution of the saccade [20]. Pollatsek and colleagues measured the perceptual span in bilingual Israeli readers who spoke English as their second language. They found that bilingual Israeli readers showed an asymmetry perceptual span that extended 14 characters to the left of fixation and 4 characters to the right while reading Hebrew [55]. Additionally, although no overall extent of the perceptual span was examined, Jorden et al., reported a leftward asymmetry in perceptual span when participants read Arabic [56]. McConkie and Rayner reported that skilled readers of English and other alphabetic languages reading from left to right showed an asymmetric perceptual span, extending 14-15 characteristics to the right of fixation and 3-4 characteristics to the left [57]. In contrast, Chinese readers showed a narrower perceptual span that extended 1 character space leftward and 3 characters spaces rightward as reported in Inhoff and Liu [58] or extended beyond 4 characters spaces rightward depending on the font size as reported in Yan et al. [59]. It is possible that the early disengagement of attention in Chinese participants leads to a reduction in prosaccade latency because the 5^°^targets in the present study exceed the perceptual span used to acquire useful information by Chinese participants (the size of each character in [58] study was 0.9°), but still fall into the perceptual span of the Arabic and Persian participants. Accordingly, the Arabic and Persian group showed a directional difference of prosaccade latency whereas no group difference was found for the 10^°^target away from the centre.

### 4.2 Impact of habitual reading direction on prosaccade gain

Inconsistent with other studies [44, 60, 61], we did not find a significant gain difference with respect to stimulus magnitude. One possible explanation could be the different perceptual span of the 2 groups of the participants. Therefore, hypometric performance towards a 5^°^target disappeared when we combined the 2 groups together. A significant interaction between group and stimulus direction was found in prosaccade gain. Although Arabic and Persian subjects made more accurate prosaccade towards the side of their non-habitual reading direction (i.e., right side of the fixation), Chinese participants also performed similarly that target at the right side elicited more accurate prosaccades. This finding was consistent with the result as reported in (Vergilino-Perez et al., 2012) that rightward prosaccade had larger amplitude. Nevertheless, no primary habitual reading direction effect was found on prosaccade gain.

### 4.3 Impact of habitual reading direction on antisaccade latency

The current result agreed with previous findings where the latency of a correct horizontal antisaccade was independent of stimulus direction or magnitude [44, 62, 63]. Although different eye movement paradigms (overlap and gap conditions) were tested on different participant cohorts, the latency of the correct antisaccades did not show the same pattern as the prosaccades. The study reported by Knox et al. also revealed that the correct antisaccade latency was identical between Chinese participants who exhibited a high proportion of express saccades and those who did not [52]. Reading relies more on perceptually driven saccades [64]. In contrast, cognition is needed to inhibit the reflexive error that was stimulated by a perceptual stimulus in the antisaccade task [8]. Therefore, the possibility that the cognitive difference induced by subjects’ habitual reading habits had greater influence on reflexive saccades compared to volitional saccades. That is, reaction time of a reflexive saccade was more impacted by reading direction than initiation time of an antisaccade. Therefore, the correct antisaccade latency was not significantly affected by the habitual reading direction or the direction of the stimulus.

### 4.4 Impact of habitual reading direction on antisaccade error rate

The antisaccade error rate was found to be significantly higher in the Chinese group compared with the Arabic and Persian group for the 10^°^with-direction stimulus presentation, although shorter prosaccade latency was observed in the Chinese group when the target appeared 5^°^along their habitual reading direction. Previous study showed that prosaccade training increased the number of antisaccade errors, because the reinforcement of the practice made it harder to inhibit a reflexive glance. Accordingly, subjects who were trained on antisaccade eye movement significantly reduced their directional errors [65]. The Arabic and Persian participants in the present study were familiar with reading in both directions, the inhibition of reading in the opposite direction during one language processing might improve their ability to supress reflexive saccades, which resulted in significantly fewer number of errors. Although Arabic and Persian participants had shorter prosaccade latency towards their non-habitual reading direction side, they did not show higher antisaccade error rate, or more antisaccade errors towards the right side. This implies that this quicker prosaccade latency was not fast enough to make more antisaccade errors.

### 4.5 Impact of habitual reading direction on self-paced saccades

The self-paced saccade task has been considered as an almost entirely volitional eye movement task as no reflexive cues are presented to trigger saccadic eye movements [9]. The generation of self-paced saccades requires a series of quick volitional engagements and disengagements of attention between 2 static stimuli. Although it has been proposed that language processing drives the disengagement and shift of attention to the next word of interest in the direction of reading [66], the current result failed to find a difference in the mean inter-saccadic interval between the self-paced saccades initiated to the side of subjects’ habitual reading direction and those to the non-habitual reading direction in both groups. One possible explanation is that the amplitude of the saccades required to execute the self-paced saccade task is much larger than those produced during normal reading, thus the difference was not shown in the current ocular motor task. Alternatively, it is possible that the subjects’ sustained task engagement was more relevant to the performance of the self-paced saccade task, compared to the attentional modulation, as a result of the need to continuously initiate and execute eye movements [67]. Therefore, inter-saccadic interval was not significantly different between groups. However, both groups’ participants showed more accurate gain when they made a self-paced saccade towards the right target. This result was similar with the prosaccades, where participants had more accurate gain when the target appeared at the right side.

### 4.6 Limitations of the study

At present, very few studies have investigated the differences in saccadic eye movements in response to a low-cognitive demand target between populations or individuals from different cultural backgrounds. The analysis of this study has been primarily concentrated on the effect of the habitual reading direction on the directionality of saccadic eye movements. However, only the Arabic and Persian subjects who made prosaccade to a 5^°^target supported our hypothesis, Nevertheless, we would like to point out several limitations in the current experiment. First, we did not recruit monolingual Arabic or Persian participants. The lack of monolingual subjects who only read from right to left leads to an uncertainty about the impact of habitual reading direction on saccadic eye movements as the Arabic and Persian participants in the present study were experienced in reading both directions. Secondly, we did not recruit participants in addition to Chinese who habitually read from left to right (such as in an alphabetic language such as English). Therefore, it is difficult to examine if some differences in the current study were due to the culture or reading habit differences. Finally, the current study had a relatively small sample size, limiting the generalizability of our result. However exploratory, this study offers some insights into the neural activities of oculomotor behaviours among different cultures. Further study can be performed on a large-scale cohort.

## 5. Conclusions

In the current study, we aimed to find the effect of the primary habitual reading direction on the directionality of the characteristics of saccadic eye movements in healthy Chinese as well as Arabic and Persian participants using prosaccade, antisaccade and self-paced tasks. We hypothesised that participants showed shorter prosaccade latency and a higher antisaccade error rate when a stimulus was presented at the side of their non-habitual reading direction. Our hypotheses were partially accepted, with significantly shorter prosaccade latency found in the Arabic and Persian participants in responding to the 5^°^rightward target. The present study may contribute to the investigation of the neural mechanisms of oculomotor behaviours between populations.

## References

1. Carpenter RHS. The neural control of looking. Curr Biol. 2000;10(8):R291–3.

2. Irwin DE. Information integration across saccadic eye movements. Cogn Psychol. 1991;23(3):420–56.

3. Pretegiani E, Optican LM. Eye Movements in Parkinson’s Disease and Inherited Parkinsonian Syndromes. Front Neurol. 2017;8:592.

4. Carpenter RHS. Frontal cortex. Choosing where to look. Curr Biol. 1994;4(4):341–3.

5. Hutton SB, Ettinger U. The antisaccade task as a research tool in psychopathology: a critical review. Psychophysiology. 2006;43(3):302–13.

6. Massen C. Parallel programming of exogenous and endogenous components in the antisaccade task. Q J Exp Psychol A. 2004;57(3):475–98.

7. Munoz DP, Everling S. Look away: the anti-saccade task and the voluntary control of eye movement. Nat Rev Neurosci. 2004;5(3):218–28.

8. Hutton SB. Cognitive control of saccadic eye movements. Brain Cogn. 2008;68(3):327–40.

9. Abel LA, Douglas J. Effects of age on latency and error generation in internally mediated saccades. Neurobiol Aging. 2007;28(4):627–37

10. Bittencourt J, Velasques B, Teixeira S, Basile LF, Salles JI, Nardi AE, et al. Saccadic eye movement applications for psychiatric disorders. Neuropsychiatr Dis Treat. 2013;9:1393–409.

11. Broerse A, Crawford TJ, den Boer JA. Parsing cognition in schizophrenia using saccadic eye movements: a selective overview. Neuropsychologia. 2001;39(7):742–56.

12. Barkley RA. Behavioral inhibition, sustained attention, and executive functions: constructing a unifying theory of ADHD. Psychol Bull. 1997;121(1):65–94.

13. Gould TD, Bastain TM, Israel ME, Hommer DW, Castellanos FX. Altered performance on an ocular fixation task in attention-deficit/hyperactivity disorder. Biol Psychiatry. 2001;50(8):633–5.

14. Feifel D, Farber RH, Clementz BA, Perry W, Anllo-Vento L. Inhibitory deficits in ocular motor behavior in adults with attention-deficit/hyperactivity disorder. Biol Psychiatry. 2004;56(5):333–9.

15. Lezak MD. Neuropsychological assessment: Oxford University Press, USA; 2004.

16. Lemos J, Pereira D, Almendra L, Rebelo D, Patrício M, Castelhano J, et al. Distinct functional properties of the vertical and horizontal saccadic network in Health and Parkinson’s disease: An eye-tracking and fMRI study. Brain Res. 2016;1648(Pt A):469–84.

17. Winograd-Gurvich C, Georgiou-Karistianis N, Fitzgerald PB, Millist L, White OB. Ocular motor differences between melancholic and non-melancholic depression. J Affect Disord. 2006;93(1-3):193–203.

18. Fischer B, Weber H. Effects of stimulus conditions on the performance of antisaccades in man. Exp Brain Res. 1997;116(2):191–200.

19. Liversedge SP, Findlay JM. Saccadic eye movements and cognition. Trends Cogn Sci. 2000;4(1):6–14.

20. Hoffman JE, Subramaniam B. The role of visual attention in saccadic eye movements. Percept Psychophys. 1995;57(6):787–95.

21. Roberts RJ, Hager LD, Heron C. Prefrontal cognitive processes: Working memory and inhibition in the antisaccade task. J Exp Psychol Gen. 1994;123(4):374.

22. Dick S, Kathmann N, Ostendorf F, Ploner CJ. Differential effects of target probability on saccade latencies in gap and warning tasks. Exp Brain Res. 2005;164(4):458–63.

23. Saslow MG. Effects of components of displacement-step stimuli upon latency for saccadic eye movement. J Opt Soc Am. 1967;57(8):1024–9.

24. Carpenter RHS, Williams ML. Neural computation of log likelihood in control of saccadic eye movements. Nature. 1995;377(6544):59–62.

25. Inhoff AW, Pollatsek A, Posner MI, Rayner K. Covert attention and eye movements during reading. Q J Exp Psychol A. 1989;41(1):63–89.

26. Han S, Northoff G. Culture-sensitive neural substrates of human cognition: A transcultural neuroimaging approach. Nat Rev Neurosci. 2008;9(8):646.

27. Han S, Northoff G. Reading direction and culture. Nat Rev Neurosci. 2008;9(12):965.

28. Vaid J, Singh M. Asymmetries in the perception of facial affect: is there an influence of reading habits? Neuropsychologia. 1989;27(10):1277–87.

29. Chokron S, De Agostini M. Reading habits influence aesthetic preference. Brain Res Cogn Brain Res. 2000;10(1-2):45–9.

30. Chokron S, Imbert M. Influence of reading habits on line bisection. Brain Res Cogn Brain Res. 1993;1(4):219–22.

31. Eviatar Z. Reading direction and attention: effects on lateralized ignoring. Brain Cogn. 1995;29(2):137–50.

32. Kazandjian S, Chokron S. Paying attention to reading direction. Nat Rev Neurosci. 2008;9(12):965–5.

33. Rinaldi L, Di Luca S, Henik A, Girelli L. Reading direction shifts visuospatial attention: an Interactive Account of attentional biases. Acta Psychol (Amst). 2014;151:98–105.

34. Afsari Z, Ossandón JP, König P. The dynamic effect of reading direction habit on spatial asymmetry of image perception. J Vis. 2016;16(11):8.

35. Rayner K. Eye guidance in reading: fixation locations within words. Perception. 1979;8(1):21–30.

36. Nuthmann A, Engbert R, Kliegl R. Mislocated fixations during reading and the inverted optimal viewing position effect. Vision Res. 2005;45(17):2201–17.

37. Deutsch A, Rayner K. Initial fixation location effects in reading Hebrew words. Lang Cogn Process. 1999;14(4):393–421.

38. Yan M, Zhou W, Shu H, Yusupu R, Miao D, Krügel A, et al. Eye movements guided by morphological structure: evidence from the Uighur language. Cognition. 2014;132(2):181–215.

39. Yan M, Pan J, Chang W, Kliegl R. Read sideways or not: Vertical saccade advantage in sentence reading. Read Writ. 2019;32(8):1911–26.

40. Beydagi H, Yilmaz A, Süer C. The effect of direction on saccadic eye movement parameters. J Basic Clin Physiol Pharmacol. 1999;10(1):73–7.

41. Constantinidis TS, Smyrnis N, Evdokimidis I, Stefanis NC, Avramopoulos D, Giouzelis I, et al. Effects of direction on saccadic performance in relation to lateral preferences. Exp Brain Res. 2003;150(4):443–8.

42. Tagu J, Doré-Mazars K, Lemoine-Lardennois C, Vergilino-Perez D. How Eye Dominance Strength Modulates the Influence of a Distractor on Saccade Accuracy. Invest Ophthalmol Vis Sci. 2016;57(2):534–43.

43. Killgore WD, Yurgelun-Todd DA. Activation of the amygdala and anterior cingulate during nonconscious processing of sad versus happy faces. Neuroimage. 2004;21(4):1215–23.

44. Vergilino-Perez D, Fayel A, Lemoine C, Senot P, Vergne J, Doré-Mazars K. Are there any left-right asymmetries in saccade parameters? Examination of latency, gain, and peak velocity. Invest Ophthalmol Vis Sci. 2012;53(7):3340–8.

45. Irving EL, Steinbach MJ, Lillakas L, Babu RJ, Hutchings N. Horizontal saccade dynamics across the human life span. Invest Ophthalmol Vis Sci. 2006;47(6):2478–84.

46. Fischer B, Breitmeyer B. Mechanisms of visual attention revealed by saccadic eye movements. Neuropsychologia. 1987;25(1A):73–83.

47. Fischer B. The preparation of visually guided saccades. Rev Physiol Biochem Pharmacol. 1987;106:1–35.

48. Fischer B, Ramsperger E. Human express saccades: extremely short reaction times of goal directed eye movements. Exp Brain Res. 1984;57(1):191–5.

49. De Clerck M, Crevits L, Van Maele G. Saccades: is there a difference between right and left? Neuroophthalmology. 2000;24(2):327–30.

50. Amatya N, Gong Q, Knox PC. Differing proportions of ‘express saccade makers’ in different human populations. Exp Brain Res. 2011;210(1):117–29.

51. Knox PC, Wolohan FD. Cultural diversity and saccade similarities: culture does not explain saccade latency differences between Chinese and Caucasian participants. PLoS One. 2014;9(4):e94424

52. Knox PC, Amatya N, Jiang X, Gong Q. Performance deficits in a voluntary saccade task in Chinese “express saccade makers”. PLoS One. 2012;7(10):e47688.

53. Carpenter RHS. Oculomotor procrastination. Eye movements: cognition and visual perception. Fischer DF, Monty RA, eds. 1981. pp 237–46.

54. Carpenter RHS. Express saccades: is bimodality a result of the order of stimulus presentation? Vision Res. 2001;41(9):1145–51.

55. Pollatsek A, Bolozky S, Well AD, Rayner K. Asymmetries in the perceptual span for Israeli readers. Brain Lang. 1981;14(1):174–80.

56. Jordan TR, Almabruk AAA, Gadalla EA, McGowan VA, White SJ, Abedipour L, et al. Reading direction and the central perceptual span: Evidence from Arabic and English. Psychon Bull Rev. 2014;21(2):505–11.

57. McConkie GW, Rayner K. The span of the effective stimulus during a fixation in reading. Percept Psychophys. 1975;17(6):578–86.

58. Inhoff AW, Liu W. The perceptual span and oculomotor activity during the reading of Chinese sentences. J Exp Psychol Hum Percept Perform. 1998;24(1):20–34.

59. Yan M, Zhou W, Shu H, Kliegl R. Perceptual span depends on font size during the reading of Chinese sentences. J Exp Psychol Learn Mem Cogn. 2015;41(1):209–19.

60. Ploner CJ, Ostendorf F, Dick S. Target size modulates saccadic eye movements in humans. Behav Neurosci. 2004;118(1):237–42.

61. Kapoula Z. Evidence for a range effect in the saccadic system. Vision Res. 1985;25(8):1155–7.

62. Bell AH, Everling S, Munoz DP. Influence of stimulus eccentricity and direction on characteristics of pro- and antisaccades in non-human primates. J Neurophysiol. 2000;84(5):2595–604.

63. Fischer B, Biscaldi M, Gezeck S. On the development of voluntary and reflexive components in human saccade generation. Brain Res. 1997;754(1):285–97.

64. Feng G. Is there a common control mechanism for anti-saccades and reading eye movements? Evidence from distributional analyses. Vision Res. 2012;57:35–50.

65. Dyckman KA, McDowell JE. Behavioral plasticity of antisaccade performance following daily practice. Exp Brain Res. 2005;162(1):63–9.

66. Morrison RE. Manipulation of stimulus onset delay in reading: evidence for parallel programming of saccades. J Exp Psychol Hum Percept Perform. 1984;10(5):667.

67. Azizi E. The influence of playing video games as an attention rehabilitation technique in patients with traumatic brain injury. Published PhD thesis. Unoversity of Melbourne. 2016.

